# Repurposing the orphan drug nitisinone to control the transmission of African trypanosomiasis

**DOI:** 10.1101/2020.06.08.139808

**Authors:** Marcos Sterkel, Lee R. Haines, Aitor Casas-Sánchez, Vincent Owino Adung’a, Raquel J. Vionette-Amaral, Shannon Quek, Clair Rose, Mariana Silva dos Santos, Natalia Garcia Escude, Hanafy Ismael, Mark I. Paine, Seth M. Barribeau, Simon Wagstaff, James I. MacRae, Daniel Masiga, Laith Yakob, Pedro L. Oliveira, Álvaro Acosta-Serrano

## Abstract

Tsetse transmit African trypanosomiasis, which is a disease fatal to both humans and animals. A vaccine to protect against this disease does not exist so transmission control relies on eliminating tsetse populations. Although neurotoxic insecticides are the gold standard for insect control, they negatively impact the environment and reduce insect pollinator species. Here we present a promising, environment-friendly alternative that targets insect tyrosine metabolism pathway. A bloodmeal contains high levels of tyrosine, which is toxic to haematophagous insects if it is not degraded. RNAi silencing of either the first two enzymes in the tyrosine degradation pathway (TAT and HPPD) was lethal to tsetse. Furthermore, nitisinone (NTBC), an FDA-approved tyrosine catabolism inhibitor, killed tsetse regardless if the drug was orally or topically applied. However, it did not affect bumblebee survival. A mathematical model shows that NTBC could reduce the transmission of African trypanosomiasis in sub-Saharan Africa, thus accelerating current elimination programmes.

## Introduction

Human African trypanosomiasis (HAT), also known as sleeping sickness, is a parasitic disease caused predominantly by the parasite *Trypanosoma brucei gambiense*. These parasites are transmitted when infected tsetse flies (*Glossina spp*.) blood feed. HAT currently affects 3,500 people/year; most patients live in the Democratic Republic of the Congo and an estimated 70 million people remain at risk of infection in sub-Saharan Africa^1^. Tsetse also spread animal African trypanosomiasis (AAT), which causes high mortality rates in livestock and consequently severely limits animal production^2^. As no vaccine for either HAT or AAT exist, and efficacious treatments are often difficult to obtain, tsetse population control remains essential to limit the spread of trypanosomiasis. In the last decades, tsetse control tools such as aerial spraying of insecticides (pyrethroids), visual and odour baited traps, insecticide-treated livestock, live traps, insecticide-impregnated traps and targets, and sterile male releases have been employed^3–7^. Despite such efforts, AAT and HAT persist and both economic development and public health continue to be jeopardised^8^. Consequently, a novel complementary strategy to control these parasitic diseases is highly desired. Tsetse, like other blood-feeding arthropods, ingest large quantities of blood and often exceed twice their body weight in a single meal^9^. Since more than 80% of blood dry weight consists of proteins, large quantities of amino acids are released in the midgut during bloodmeal digestion^10^. Previously, we showed that blocking tyrosine catabolism after a bloodmeal is lethal in mosquitoes, ticks and kissing bugs due to the accumulation of toxic quantities of tyrosine^11^. However, inhibiting tyrosine catabolism in non blood-feeding insects is harmless, which further provides evidence for the essentiality of this pathway for haematophagy^11^. In the present work, we evaluated how tsetse physiology was controlled by two enzymes in the tyrosine catabolism pathway: tyrosine aminotransferase (TAT) and 4-hydroxyphenylpyruvate dioxygenase (HPPD). The drug nitisinone (NTBC) is an HPPD inhibitor currently used to treat patients suffering from the genetic disease tyrosinemia type I^12^ and is under clinical evaluation for the treatment of alkaptonuria^13^. NTBC was lethal to blood-fed tsetse flies. NTBC treatment, either administered orally as an endectocide or topically, causes the accumulation of tyrosine and 4-hydroxyphenyl lactic acid (HPLA) metabolites, which leads to initial fly paralysis followed by tissue destruction within 18 hours of the bloodmeal. Our results provide evidence that NTBC could be used as an eco-friendly synergistic strategy alongside current tsetse control practices.

## Results

### Tyrosine detoxification is essential in tsetse

Tyrosine catabolism is a highly conserved pathway (Fig. 1A) in most eukaryote and prokaryote species with only a few exceptions such as the pea aphid, *Acyrthosiphon pisum*^14^ and trypanosome parasites^15^. The genes encoding *TAT* and *HPPD* were identified in five *Glossina* species^16^, as well as in all hematophagous arthropod species with sequenced genomes (Supplementary Figure 1). Transcript knockdown using RNA interference, of either *TAT* or *HPPD* genes, was lethal to flies once they fed on bloodmeal (Fig. 1B and Supplementary Figure 2). This lethality was further validated by feeding flies with blood spiked with mesotrione, an HPPD inhibitor widely used as a selective herbicide on corn crops under the brand name *Callistro*^®^, Syngenta (Supplementary Figure 3A). No differences in susceptibility to mesotrione were observed between fly sex or age (data not shown). The mesotrione concentration that killed 50% of the insects 24 hours after administration (LC_50_) was 357.7 μM (CI: 222.5-512.4) (Supplementary Figures 3A and 3C). This concentration is ~30X higher than the drug concentration detected in human plasma (4 μg/ml (11.78 μM)) after volunteers received an oral dose of 4 mg/kg body weight^17^.

**Fig. 1.**
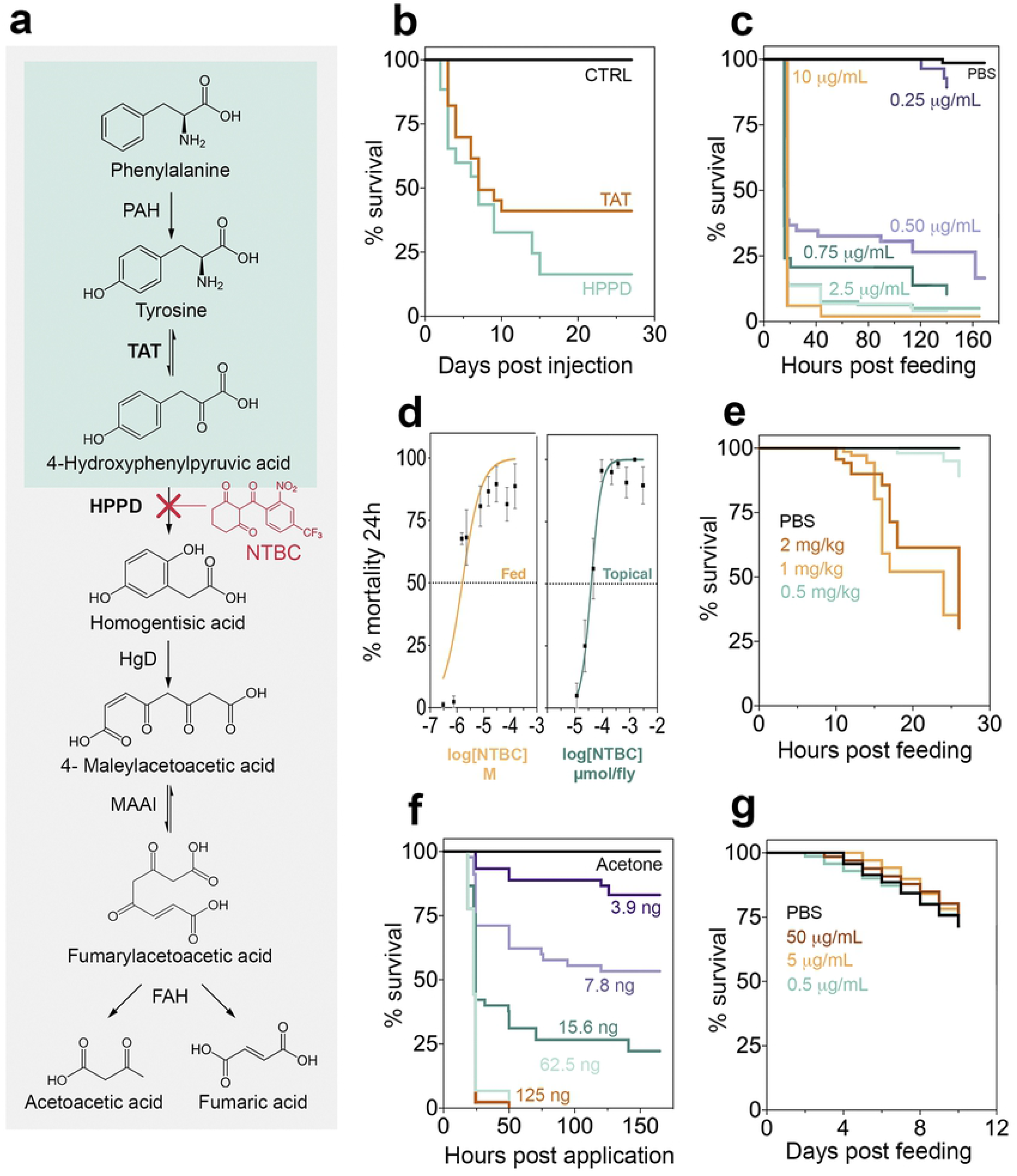
Inhibiting tyrosine catabolism is lethal for tsetse, but not for bees. **a** Tyrosine catabolism pathway. PAH: Phenylalanine hydroxylase; HgD: Homogentisate 1,2-dioxygenase; MAAI: Maleylacetoacetate isomerase; FAH: Fumarylacetoacetase. **b** Survival of *G. m. morsitans* injected with dsRNA for TAT and HPPD. Two independent experiments were performed, each with n=10-13, 12-16 and 12-14 for dsGFP (control-CTRL), dsTAT and dsHPPD, respectively. Five insects of each group were dissected on day 3 post-bloodmeal (PBM) to assess gene knockdown efficiency (Supplementary figure 2). **c** Survival of *G. m. morsitans* fed with horse blood supplemented with NTBC or PBS (Control. 9:1; v/v). Three to six independent experiments were performed for different doses, each with n =10-25 insects. **d** Dose-response curves for *G. m. morsitans* survival 24 hours PBM (Post-bloodmeal): NTBC feeding and topical application assays (Media ± SEM. All doses applied are shown in Supplementary figure 3B). **e** Male *G. pallidipes* survival after feeding on PBS- or NTBC-treated rats. Doses are presented as mg of NTBC per kg of rat body weight. Three independent experiment were performed for different doses, each with n=20-40. **f** Survival of NTBC topically treated *G. m. morsitans* after a bloodmeal. Three independent experiments were performed for different doses, each with n=10-20 insects. **g** Bees survival maintained with 50% of sugar solution supplemented with NTBC (or PBS) and fed with pollen. Two independent experiments were performed, each with n=25 insects.

### NTBC ingestion is lethal to tsetse

Hypertyrosinemia Type I (HT-1) is a severe human genetic disease caused by a mutation in the gene encoding for the last enzyme of the tyrosine catabolism pathway, fumarylacetoacetase (FAH). This mutation causes the accumulation of toxic metabolites in blood and tissues. The only drug available to minimise the effect of HT-1 is the orphan drug nitisinone (NTBC / Orfadin^®^). NTBC is an HPPD inhibitor that prevents a build-up of toxic products derived from fumarylacetoacetate accumulation^18^. NTBC is remarkably safe to use with few reported side effects in <1% of patients^19^. When NTBC was artificially fed to tsetse flies in the bloodmeal, it was ~173 times more potent than mesotrione with a LC_50_= 2.07 μM (CI: 0.709 to 4.136) (Fig. 1C and Fig. 1D). This lethal concentration is ~12 times lower than the concentration of NTBC persisting in human plasma (8 μg/ml (24.3 μM)) after administering a standard therapeutic oral dose (1 mg/kg). The NTBC half-life in human plasma is 54 hours^17^, so assuming linear drug degradation, human blood would remain toxic to tsetse up to ~7 days after a single dose in the therapeutic range. These results suggest that NTBC used as endectocide (here defined as a drug or insecticide that is administered to humans or animals to control parasites) could be used to rapidly decrease tsetse populations as part of a drug-based vector control strategy. Furthermore, it did not matter if the tsetse flies were infected with *T. brucei*; NTBC lethality remained unaltered in infected flies (Supplementary Figure 4).

### Tsetse fed on NTBC-treated mice die

To evaluate the potential of using NTBC as an endectocide, the *in vivo* efficacy of NTBC was assessed. Colony-reared tsetse (*G. pallidipes)* were allowed to feed on rats that had been orally treated with different NTBC concentrations. We observed that after feeding on rats receiving an oral dose of NTBC equal to 1 mg/kg (the therapeutic dose for humans with Tyrosinemia type I) killed *G. pallidipes* as reflected in our previous data (Fig. 1E). These data provide direct evidence that NTBC could be used as an endectocide for tsetse control.

### NTBC causes insect paralysis and systemic tissue destruction

Treatment of tsetse with either mesotrione- or NTBC-spiked blood presented the same unique physiological changes, which has two distinct phases we classified as early stage and death phenotype. The early-stage phenotype was observed 8-10 hours after NTBC (or mesotrione) ingestion in the bloodmeal; i.e. the flies remained alive (evidenced by red eyes) but were unable to move (often upside-down; Supplementary Video 1, Supplementary Figure 13). Exposure to bright light or tarsal stimulation often produced only a short burst of leg waving. Small, white, rhomboidal crystals (previously identified as tyrosine crystals^11^) were observed on the outside surface of the midgut epithelium against the body wall cuticle. Also, dark brown melanin-like deposits had formed in different tissues, such as the fat body, salivary glands and flight muscle (Fig. 2A and Fig. 2B). Disorganization of tissues in the digestive tract, such as the proventriculus and midgut was evident (Fig. 2C), and likely contributed to intestinal fragility, compromised digestion and leaking of the midgut content into the haemocoel.

**Fig. 2.**
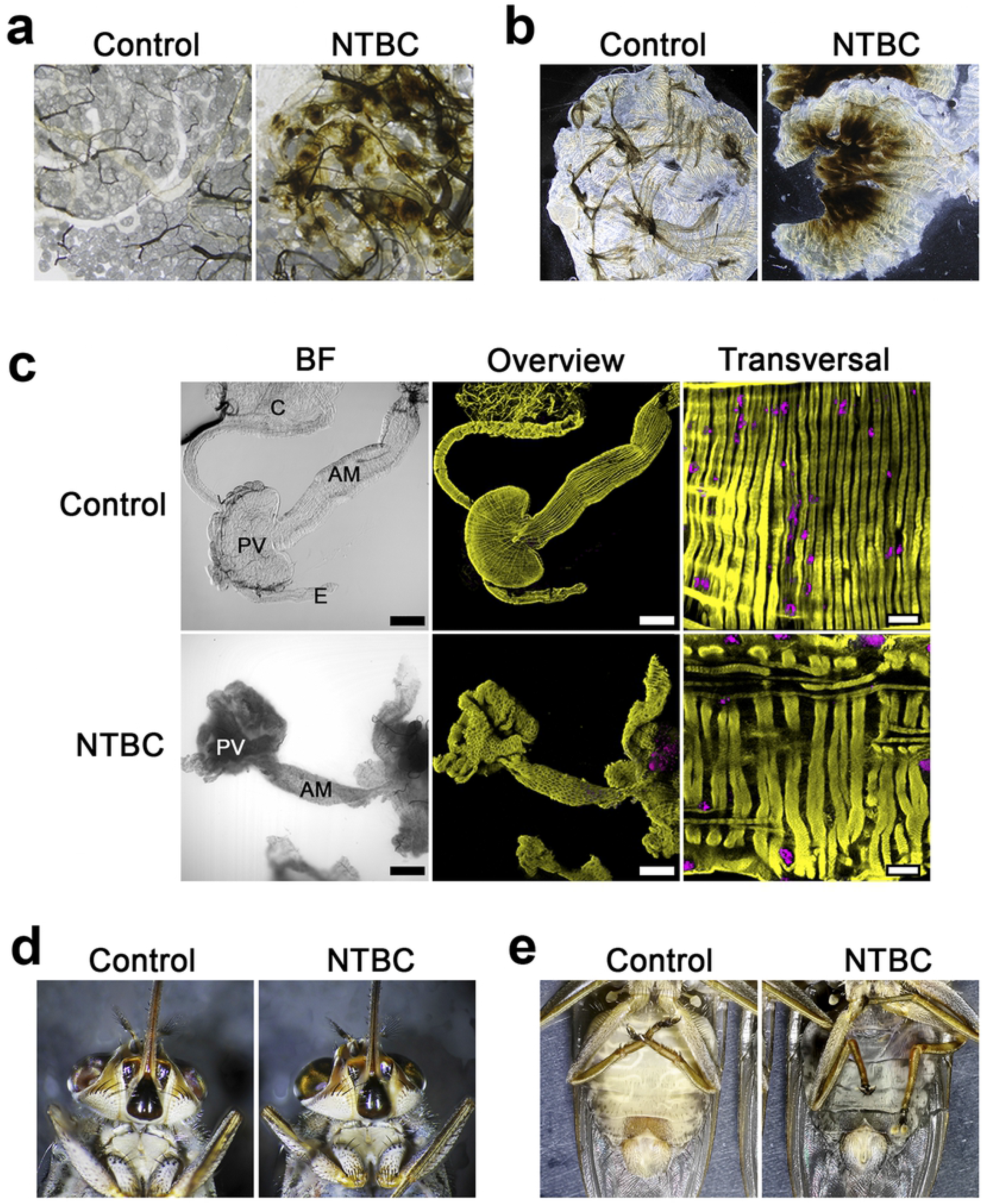
Tsetse phenotypes observed upon NTBC treatment. **a** Fat Body. **b** Flight muscle. **c** Bright field (left) and fluorescence 3D-reconstructed (middle) overviews of the anterior midgut (AM), proventriculus (PV), crop (C) and oesophagus (E) from tsetse fed either without (Control) or with NTBC. Fluorescence channels merge SiR-actin staining (yellow) and DAPI (magenta). Scale bars: 200 μm (100X). Detailed sections (right) of transversal muscular fibres from the outer muscular tissues in the anterior midgut region. Scale bars=20 μm (630X). **d** and **e** Death phenotype. Flies are already dead and the abdomens are back and filled with undigested blood.

The death phenotype was observed between 15-18 hours after treatment and was characterised by a prolonged shift in eye colour from red to golden brown (Fig. 2D). Treated tsetse showed extreme cuticular thinning and complete loss of abdominal elasticity. The abdomen was strikingly distended, dark and filled with blood that had leaked into the haemocoel due to damage of the gut epithelium (Fig. 2E). The ingested blood was darkened as though it had been oxidized. Most of the internal tissues and organs (e.g. gut, testes, ovaries, salivary glands, Malpighian tubules and flight muscle) were completely destroyed and could not even be recognized in the abdomen during dissection, suggesting extensive autolysis; only the hindgut, rectum, and trachea remained identifiable. Furthermore, the fat body also disappeared and only lipid droplets could be seen floating in the haemocoel (Supplementary Figure 5). From the outside, the thorax appeared normal, but upon dissection, we observed the flight muscles had detached from the thorax and become melanised, which explains why tsetse quickly lost the ability to fly.

### NTBC leads to an accumulation of free tyrosine and 4-hydroxyphenyl lactic acid

To better understand the metabolic changes in tsetse exposed to NTBC, the haemolymph of NTBC-treated (and control) flies was collected at different time-points. Metabolomics analysis revealed increased levels of tyrosine and 4-hydroxyphenyl lactic acid (HPLA, Fig. 3). The knockdown of *TAT* gene, which is expected to reduce levels of 4-hydroxyphenylpyruvate (HPPA) and, in consequence, produce less HPLA, caused the same lethal phenotype observed upon HPPD inhibition. Together, this suggests tyrosine accumulation is the likely cause of the flies’ death. Furthermore, feeding the flies with blood spiked with HPLA or the injection of HPLA into the fly haemocoel did not affect their survival (Supplementary Table 1). In contrast to tyrosine and phenylalanine, the titre of all other detected amino acids was reduced (Fig. 3C and Fig. 3D), which may reflect a lower digestion rate in NTBC-treated flies.

**Fig. 3.**
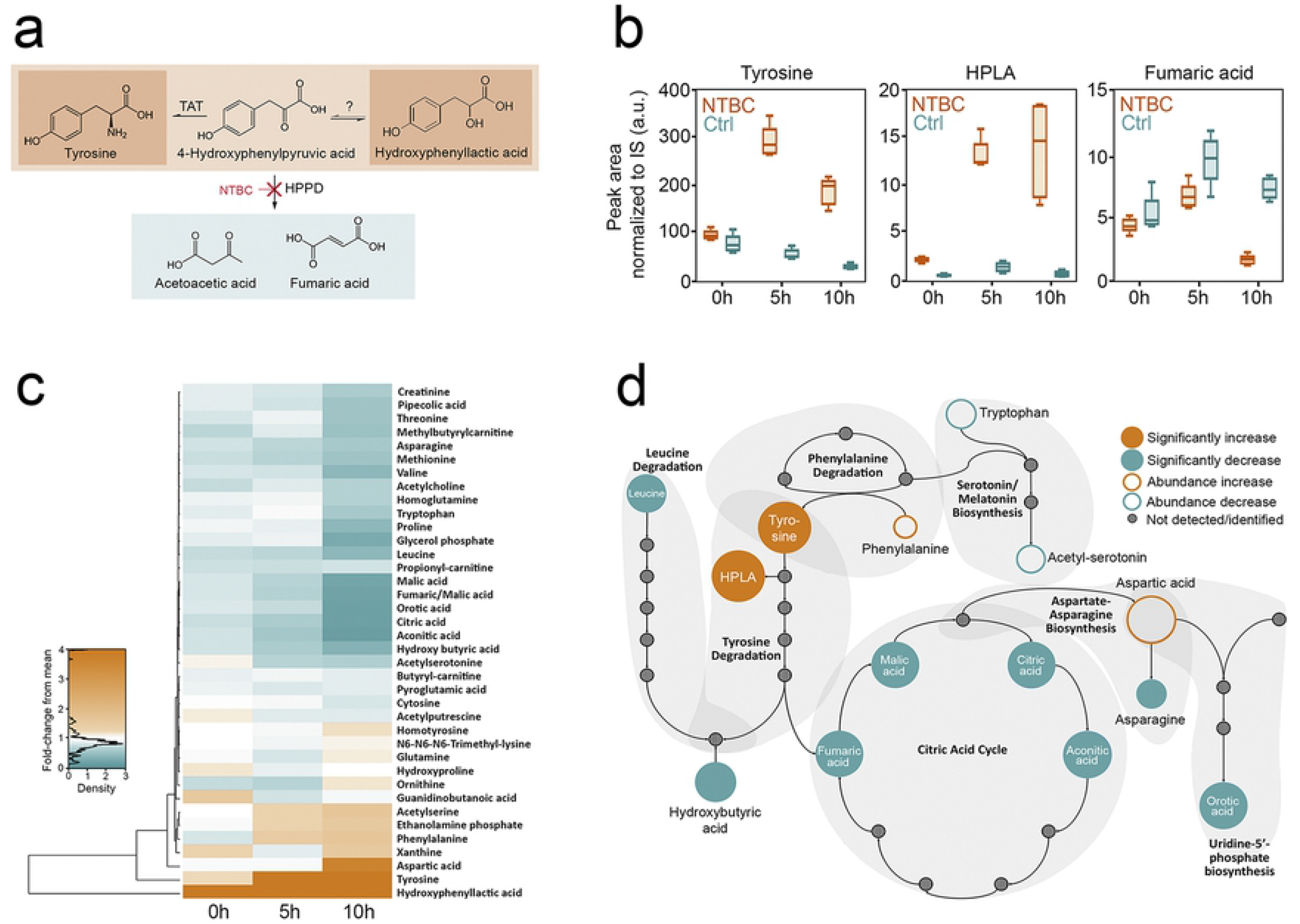
Metabolomic analysis of tsetse haemolymph upon NTBC treatment at different times post-bloodmeal (0, 5 and 10 hours). Five samples (in each group and for each time point) collected from two independent experiments were processed for and analysed by mass spectrometry **a** Schematic representation of the tyrosine catabolic pathway highlights the main metabolites accumulated and the decreased production of final products. **b** Peak area normalized to internal standards of differentially regulated metabolites in the tyrosine catabolism pathway. **c** A heat map graphically presents the metabolites identified**. d** Schematic representation of the metabolic pathways indicate metabolites that were differentially regulated in NTBC-treated flies.

### NTBC-associated damage does not depend on phenoloxidase activation or haem, but protein concentration in the bloodmeal

As extensive tissue melanisation was a result of NTBC exposure, we assessed whether systemic phenoloxidase (PO) activation led to tissues damage. PO produces melanin, but also reactive intermediates and cytotoxic quinones that could harm the flies if produced in excess^20^. To test this, we blocked PO activity by feeding tsetse on blood containing NTBC (0.01 mg/ml) and phenylthiourea (0.1 mg/ml), a PO inhibitor^21^. The addition of phenylthiourea to the bloodmeal failed to revert the lethal phenotype suggesting that PO activation is not required for NTBC death (Supplementary Figure 6).

Another potential cause of toxicity could be haem. Haem is a product of blood (haemoglobin) digestion and becomes toxic by amplifying reactive oxygen species^22^. To investigate if haem contributed to the lethal phenotype, flies were fed with horse serum (red blood cells were removed to reduce haemoglobin) supplemented with mesotrione. Fly mortality remained high even in the absence of haemoglobin, which suggested haem was not involved in the toxicity of HPPD inhibitors. Furthermore, this experiment demonstrated that the protein content in serum (around 60 mg/ml)^23^ is high enough to produce toxic levels of tyrosine (Supplementary Figure 7).

In order to investigate the toxicity of HPPD inhibitors, flies were fed with sugar supplemented with a fixed concentration of NTBC (lethal when added to bloodmeal) and different concentrations of bovine serum albumin (BSA) as the sole protein source. As expected, flies that fed on NTBC-spiked sugar (protein-free) did not die, whereas NTBC toxicity showed a clear dependence on the protein content of the meal. The observed LC_50_ was 12.5 mg/ml of protein (CI: 11.09-13.67) (Supplementary Figure 8).

### Topical application of NTBC kills tsetse

To investigate if NTBC-induced mortality was restricted to ingestion, we tested if the drug could be absorbed through the insect cuticle. Topically applying NTBC to the fly thorax caused high mortalities after tsetse took a bloodmeal, with a LD_50_ of 39 picomoles/fly (CI: 9-90) (Fig. 1D and F), 24 hours after bloodmeal ingestion. We compared this NTBC dose against a standard pyrethroid insecticide (deltamethrin) that tsetse are highly susceptible to and the LD50 was 0.116 picomoles/fly (CI: 0.093-0.145) (Supplementary Figure 9), which means that deltamethrin is 336 times more potent than NTBC.

In the field situation where HPPD inhibitors could be absorbed through the insect cuticle, fly contact with the drug could occur either before or after a bloodmeal. Thus, it is important to know how long NTBC activity persists in insects, particularly if tsetse feed after cuticular exposure. High fly mortality was observed even when a tsetse ingested a bloodmeal six days following a single topical application of NTBC (Supplementary Figure 10). To address the opposite situation where blood-feeding precedes topical exposure to NTBC, we assessed fly mortality at different times post-bloodmeal. Topically applied NTBC reduced tsetse survival up to 48 hours after a bloodmeal (Supplementary Figure 11), which implies that bloodmeal proteins were digested after two days (as predicted for tsetse) and excess tyrosine had been catabolized. In contrast to NTBC, topical application of mesotrione was not lethal, which could be due to reduced penetration through the tsetse cuticle (Supplementary Figure 12).

### NTBC is not metabolised by insect P450 enzymes

Metabolic resistance due to increased rates of insecticide metabolism by P450s can cause resistance liabilities for new compounds. However, nitisinone appears to only be moderately metabolised by CYP3A4 in humans^24^, with little oxidative metabolism by other liver CYP enzymes^25^. We have incubated nitisinone with microsomes extracted from tsetse, *Aedes* and *Anopheles* and failed to detect evidence of metabolism as measured by substrate depletion/turnover (Supplementary Table 2). Furthermore, we find no obvious metabolism by recombinant *An. gambiae* CYP6P3, a P450 with broad substrate specificity similar to CYP3A4, and associated with cross-resistance to pyrethroids and other insecticides^26^. Overall, this suggests that nitisinone may be a weak substrate for P450 metabolism in these insects. This does not preclude the evolution of metabolic resistance, but suggests it may be less likely to emerge rapidly.

### NTBC is highly stable under environmental conditions

Considering uses for NTBC in the field, we examined several stressors that could degrade drug activity and reduce its toxicity. An NTBC solution was subjected to ten freeze-thaw cycles, prolonged ambient temperature storage (5 weeks) or different exposures to light (Supplementary Table 3). These NTBC test samples (final concentration in blood 3 μM=1 μg/ml) were screened for activity by spiking them into a tsetse bloodmeal as a type of activity bioassay. In all cases, fly mortality remained unchanged, thus indicating NTBC does not quickly degrade and is stable under simple, room temperate storage conditions. In agreement with our results, Barchanska *et al*.^27^ recently demonstrated that NTBC show considerable stability under different experimental conditions such as pH of solution, temperature, time of incubation and ultraviolet radiation.

### NTBC does not affect the insect pollinator, *Bombus terrestris*

Bees are the world’s most important pollinator of food crops^28^, and several reports have shown that bee populations are directly and indirectly affected by insecticides^29^. To determine the environmental impact of NTBC on off-target species, we investigated the mortality of phytophagous pollinators when exposed to NTBC. Colony-reared *Bombus terrestris* (the buff-tailed bumblebee) were fed *ad libitum* with NTBC-spiked sugar as a sole hydration source; bee mortality was assessed over 10 days. Mortality rates did not differ between NTBC-treated and control bees during this sustained exposure, despite feeding NTBC at doses as high as 50 μg/ml (152 μM) and providing pollen as a protein source (Fig. 1G).

### A mathematical model supports use of NTBC for tsetse control

Based on our results and the pharmacokinetics of NTBC in humans^17^, we evaluated the predicted impact of using NTBC for tsetse control by mathematical modelling. Our models suggest that targeting humans in trypanosomiasis endemic areas with NTBC as an endectocide could control disease transmission with tsetse species that have a preference for feeding on humans (anthropophagic). This strategy would require drug administration with a maximal interval of every 40 days to be effective (Fig. 4A). However, tsetse flies can also feed on livestock or wildlife. Therefore, we modelled a scenario of NTBC being simultaneously given to livestock and humans. We found that tsetse population control is achieved more readily if both humans and livestock are treated, and that a frequency up to 90 days between consecutive drug administrations would still achieve control (Fig. 4B). This approach would also capture tsetse species that preferentially feed on livestock.

**Fig. 4.**
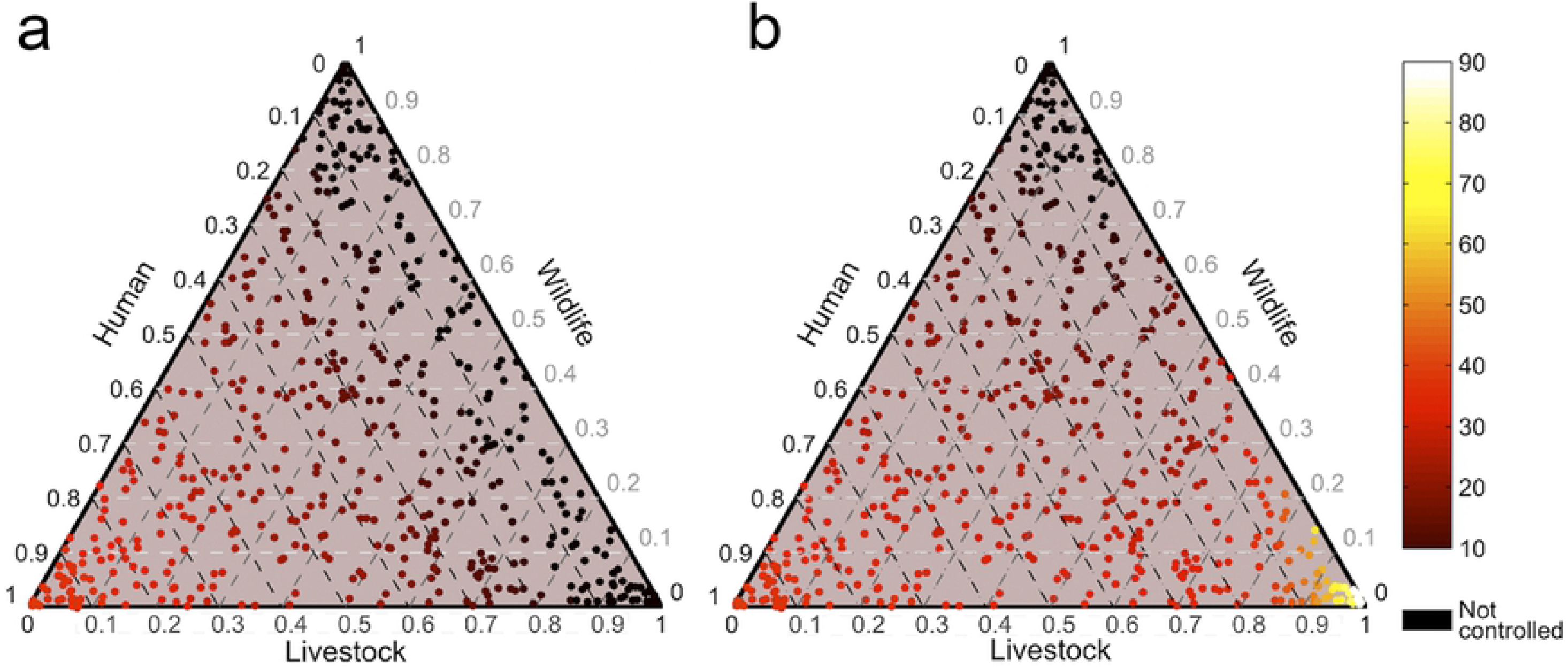
Simulation-based evaluation of the projected impact of NTBC for tsetse control. The required frequency of applying NTBC to a human population **a** and to human and livestock populations **b** is calculated to control *T. brucei* transmission. The axes denote the proportional split of bloodmeals of the local tsetse vectors between humans, livestock and wildlife. Each point (n=500) represents a random bloodmeal composition of the tsetse fly population and its colour corresponds with the frequency (in days) of application required to reduce the average basic reproduction number to below 1, as determined from a compartmental transmission model.

## Discussion

When blood-sucking insects digest a bloodmeal, large amounts of tyrosine are produced, which would be potentially toxic if it were not for the first two enzymes in the tyrosine degradation pathway, TAT and HPPD^11^. Here we provide evidence that tyrosine detoxification is essential for tsetse survival post-bloodmeal and have further investigated the possible mechanisms underlying this lethal phenotype. Furthermore, two commercially available HPPD inhibitors were evaluated as novel tsetse control interventions: mesotrione and NTBC. We concluded that NTBC is the best candidate for future field trials due to its low-dose efficacy and environmental sustainability. This drug presents unique features when compared with current insecticides as it specifically targets blood-feeding insects, has low toxicity to mammals and can be used effectively against tsetse via two delivery routes. NTBC kills tsetse upon cuticular application, similar to pyrethroids, and it can be safely administered to a mammalian host to kill tsetse upon blood-feeding.

The endectocide ivermectin has been evaluated *in vivo* as a potential tsetse control tool. *G. palpalis gambiensis* fed on cattle treated with a single ivermectin dose showed decreased survival ranging from 21-84% for the therapeutic dose (0.2 mg/kg), and from 78-94% for a dose 10 times greater (2 mg/kg)^30^. However, it is important to highlight that another tsetse species, *G. palpalis palpalis*, remained unaffected by an ivermectin dose of 2 mg/kg (4% mortality after 25 days)^31^, and no mortality was observed when flies fed on ivermectin-treated (0.5 – 1.0 mg/kg) guinea pigs and goats^32^. Similarly, no effect on fly mortality was observed when *G. tachinoides* fed on pigs treated with ivermectin^33^. These differences may be due to pharmacokinetic variations in the different host species (guinea pig, goat, cattle, and pigs) rather than *Glossina spp*-specific susceptibilities. In our study, 50% of fly mortality was observed 24 hours after tsetse took a single bloodmeal from rats that had received 1 mg/kg of NTBC. The surviving 50% were severely compromised and had lost the ability to fly. Compared to published data on ivermectin-induced fly mortality, dose-matched NTBC is more potent as it kills the flies faster than ivermectin. NTBC is also classified as a safer drug for mammals; the oral LD_50_ for NTBC in rats is >1,000 mg/kg^34^ while ivermectin is >100 times more toxic presenting a LD50=10 mg/kg^35^.

As observed in the different dose-response curves, it is not only the concentration of drug ingested or applied topically that is important, but also the quantity of blood ingested and the digestion rate of bloodmeal proteins. This is due to the particular mode of action of HPPD inhibitors. These drugs are not toxic to hematophagous arthropods during starvation or when insects are fed with low protein content diets, but their toxicity is caused by the accumulation of tyrosine derived from hydrolysis of dietary proteins. This implies that any factor affecting bloodmeal digestion, such as temperature, may also affect HPPD inhibitor toxicity and consequently how long the blood-fed arthropods take to die.

Our data shows neither the extensive melanisation observed during the early phenotype nor the haem in the bloodmeal trigger the lethal phenotype in tsetse. Similarly, in *R. prolixus* HPPD/PO and HPPD/Dopa Decarboxylase (DDC) double-gene silencing, did not rescue lethality observed in *HPPD*-silenced insects^11,36^. These similar results in tsetse reinforce the idea that PO activation and excessive melanin synthesis are not involved in killing hematophagous insects. Together with metabolomics, these results point to tyrosine accumulation (and precipitation in haemocoel and tissues) as the primary cause of insect death. Furthermore, feeding the flies with NTBC-spiked sugar supplemented with only BSA was lethal, and confirmed that tyrosine accumulation, from dietary protein digestion, caused the lethal phenotype. However, it is worth noting that the quantity of protein to be lethal upon HPPD inhibition may depend on the tyrosine content of each protein; bovine BSA tyrosine content is 3.3 % (20/608 residues). Moreover, NTBC proved to be very stable under different environmental conditions. Together, these results expand the possibility of using NTBC mixed with protein in attractive targeted sugar baits (ATSBs) to control different vector-borne diseases.

When assessing the feasibility of topically using NTBC to control tsetse flies, a high mortality was still observed 144 hours after NTBC application (Supplementary Figure 10). The longevity of this drug activity is striking and very encouraging for field-based vector control. In lab-reared colony flies, most tsetse die of dehydration/starvation after such an extensive starvation period and, of those remaining alive, many are too weak to feed when offered a bloodmeal. In wild tsetse, where fly activity and nutritional demands are greatly increased, flies are less likely to survive 144 hours of starvation so the sustained lethality of NBTC is even more encouraging. However, when comparing tsetse lethality to deltamethrin, NTBC is far less potent when topically applied to tsetse. This is not surprising as deltamethrin is a neurotoxin, while NTBC acts by a different, and albeit slower, mechanism linked to digestion and amino acid catabolism. However, the disadvantage to deltamethrin is significant; potency comes at a price. Deltamethrin is more toxic to mammals and killing is non-selective and it draws a heavy environmental penalty. All insects including pollinators and other non-target species are indiscriminately killed. Furthermore, deltamethrin can cause severe toxicity in aquatic ecosystems^37^. In contrast, NTBC toxicity profile is low (high doses are tolerated in humans and other mammals^34^) and it selectively kills blood-feeding arthropods, making it environment-friendly and more socially and ethically acceptable to employ. Despite this, pyrethroid insecticides remain the most efficacious vector control strategy for many hematophagous arthropod populations, although increasing reports of insecticide resistance compromise their efficacy^7,38–40^. Since NTBC targets a pathway separate to the neurotoxic insecticides, such as pyrethroids and ivermectin, its use could complement current vector control tools and reduce the emergence of insecticide-resistant populations.

We conclude that both NTBC and mesotrione induce rapid death in blood-fed tsetse, although the potency and the duration of their effects are very different. For these reasons, we propose using NTBC (but not mesotrione) to develop new and complementary vector control tools that target tsetse and other blood-feeding arthropod populations. The versatility of NTBC is also noteworthy. This orphan FDA approved drug could be used as a standalone technology to control tsetse and other vector populations as an endectocide, or it could complement (or provide an alternative to) the use of standard insecticides. Thus, NTBC is a safer and more environment-friendly alternative to insecticide-based vector control as it is not toxic to mammals and selectively kills only blood-feeding arthropods.

## Materials and methods

### Ethics

Male (4–6 weeks old, 180-250g in weight) Wistar rats sourced from Animal Rearing Unit and Containment Unit (ARCU) at International Centre of Insect Physiology and Ecology (*icipe*) were used. The animals were housed in standard plastic mice/rat cages (Thoren Caging Systems, Inc., Hazleton, PA, USA) with wood shavings as bedding material. The rodents were maintained on commercial pellets (Unga^®^ Kenya Ltd, Nairobi, Kenya), and water was provided *ad libitum*. Animal use and all accompanying procedures and protocols were in accordance with *The Guide for the Care and Use of Laboratory Animals* (Institute for Laboratory Animal Research. 2011). These procedures and protocols were reviewed and approved by Institutional Animal Care and Use Committee (IACUC) of *icipe* (Ref. No. IcipeACUC2018-003).

### Tsetse

The *Glossina morsitans morsitans* Westwood colony (originally from Zimbabwe) was housed in an insectary at the Liverpool School of Tropical Medicine. Flies were kept at 26°C and 73 ± 5 % relative humidity and a 12 hour light:dark cycle. The colony was maintained on defibrinated horse blood (TCS Biosciences Ltd., Buckingham, UK) and fed every 2-3 days using artificial silicone feeding membranes. Male and female experimental flies of different ages were housed and fed separately. The *G. pallidipes* (originally from Nguruman, Kenya) from Insectary Unit at *icipe* were maintained at 26°C and 75 ± 4 % relative humidity with 12 hour light:dark cycle. The flies were fed on defibrinated bovine blood using an *in vitro* silicon membrane system. Only teneral *G. pallidipes* were fed on NTBC-treated rats.

### Bumblebee rearing

Hives of *Bombus terrestris audax* ordered from Agralan (Wiltshire, UK) were kept at 27°C under constant red light and fed *ad libitum* with pollen and 50% sugar water (Ambrosia syrup, EH Thorne^®^). Only worker bees were used in this study. Five bees from five different colonies were placed into an acrylic box in five biological replicates, totalling 25 bumblebees per each experimental group. NTBC was diluted in sugar solution and placed into the box 24h after the bees acclimatized to laboratory conditions. Bees were allowed to drink *ad libitum* for 10 days. Each group received PBS or a dose of NTBC at concentrations shown to be lethal to tsetse (0.05 mg/ml, 0.005 mg/ml or 0.0005 mg/ml). Solutions were changed every five days. Worker bees were also provided with pollen (protein source) according to their rate of consumption. Bee mortality was scored daily for 10 days.

### Phylogenetic analysis

Dendograms showing the phylogenetic relationships of TAT and HPPD among insects were created according to the maximum likelihood method. Confidence values for each branch was determined through bootstrapping at 100. Analysis was performed on the web-based interface on Phylogeny.fr^41,42^, using T-Coffee for alignments under the BLOSUM62 scoring matrix and optimised with Gblocks. Dendograms were constructed with PhyML 3.0^43^. The TAT and HPPD protein sequences used in the phylogenetic analyses were taken from either Vectorbase or NCBI using the most recent gene sets available.

### Synthesis of double-stranded RNA (dsRNA)

Specific primers for *G. m. morsitans* tyrosine aminotransferase (TAT) and 4-hydroxyphenylpyruvate dioxygenase (HPPD) genes were designed using primer-blast software (NCBI. https://www.ncbi.nlm.nih.gov/tools/primer-blast) (Table 1). These primers contained the T7 polymerase binding sequence, required for dsRNA synthesis at 5’ end. The green fluorescence protein (GFP) gene amplified from the peGFP-N1 plasmid (NCBI ID: U55762.1) was used as a control dsRNA to assess *in vivo* off-target effects. PCR products were sequenced to confirm identity. MEGAscript High Yield T7 Transcription kit (Ambion) was used according to the manufacturer’s instructions to synthesize dsRNA. After synthesis, dsRNAs were precipitated by adding an equal volume of ice-cold isopropanol, and the resulting pellets were washed with ethanol, air dried and then the pellet resuspended in ultrapure water. The dsRNAs concentrations were determined spectrophotometrically at 260 nm on a NanoVue^™^ Plus Spectrophotometer (GE Healthcare, Buckinghamshire, UK) and visualized in an agarose gel (1.5% w/v) to confer dsRNA size, integrity and purity. A speedvac concentrator (ISS110. Thermo Scientific) was used to dry the dsRNAs and they were adjusted to a final concentration of 5 μg/μl in sterile nuclease-free water (NFW).

**Table 1.**
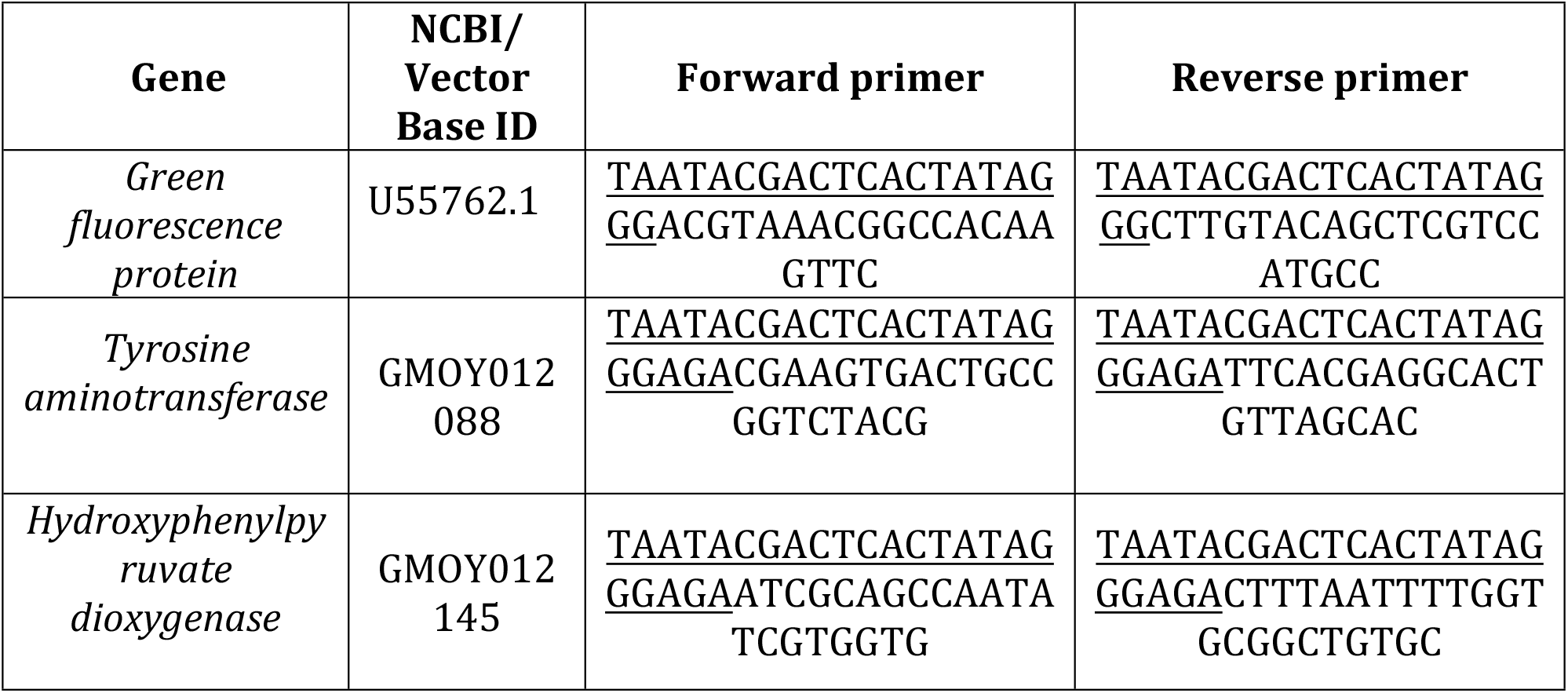
Sequences of the primers used to amplify target genes for RNAi experiments. T7 promoter sequences that were necessary for transcription are underlined.

### Gene silencing

The protocols described by Walshe *et al*.^44^ were followed for RNAi knockdown. Briefly, *G. m. morsitans* were injected in the thorax with 10 μg of each target gene dsRNA (2 μl of 5 μg/μl stock). Control insects were injected with 10 μg of GFP dsRNA. Injection needles were handmade using pulled micro-haematocrit glass capillary tubes (2.00 mm outside diameter) (Globe Scientific) mounted inside 200 μl yellow micropipette tips and sealed with Araldite epoxy glue. Flies were injected 24 hours after receiving a bloodmeal to increase survival rates. Flies were chilled for 10 minutes on ice to immobilize them and 2 μl of dsRNA was injected into the dorsolateral surface of the thorax. Injected flies were allowed to rest for 24 hours before the next bloodmeal. Following injections, flies were fed every 2-3 days on sterile, defibrinated horse blood. Three days after dsRNA injection, the digestive systems were excised to determine the efficacy of gene-silencing as evaluated by quantitative RT-PCR.

### RNA isolation and cDNA synthesis

*G. m. morsitans* flies were immobilized on ice and the digestive system was dissected into ice-cold PBS. The total RNA was extracted using TRIzol reagent (Invitrogen), according to the manufacturer’s instructions. Following treatment of the extracted mRNA with TURBO DNase (Ambion), first-strand cDNA synthesis was performed using 1 μg total RNA with “Superscript III First-strand Synthesis System for RT-PCR Kit” (Invitrogen) and poly-T primer, according to the manufacturer instructions. The cDNAs were stored at −80°C until use.

### Quantitative polymerase chain reaction (qPCR)

Specific gene-specific qPCR primers were designed to amplify a different region from that amplified by the RNAi primers to prevent dsRNA amplification. They were designed to span different exons so to easily detect contaminating DNA amplification and their efficiency was experimentally tested (Table 2). *Glossina α-tubulin* and *β-tubulin* genes were used as references (housekeeping) genes. Quantitative PCR was performed using a MxPro – Mx3005P thermocycler (Agilent Technologies) with Brilliant III Ultra-Fast SYBR^®^ Green QPCR Master Mix (Agilent Technologies) under the following conditions: 95 °C for 15 minutes, followed by 40 cycles of 95 °C for 15 seconds, 60 °C for 30 seconds and 72 °C for 30 seconds and a final extension of 72 °C for 10 minute. Target gene expression levels were assessed as 2e^-ΔCT^ values (ΔC_T_ = C_T_ gene of interest – C_T_ housekeeping gene) were used to evaluate the expression levels of the genes in the different experimental groups. (dsTAT- or dsHPPD-injected flies and dsGFP injected flies) ^45^.

**Table 2.**
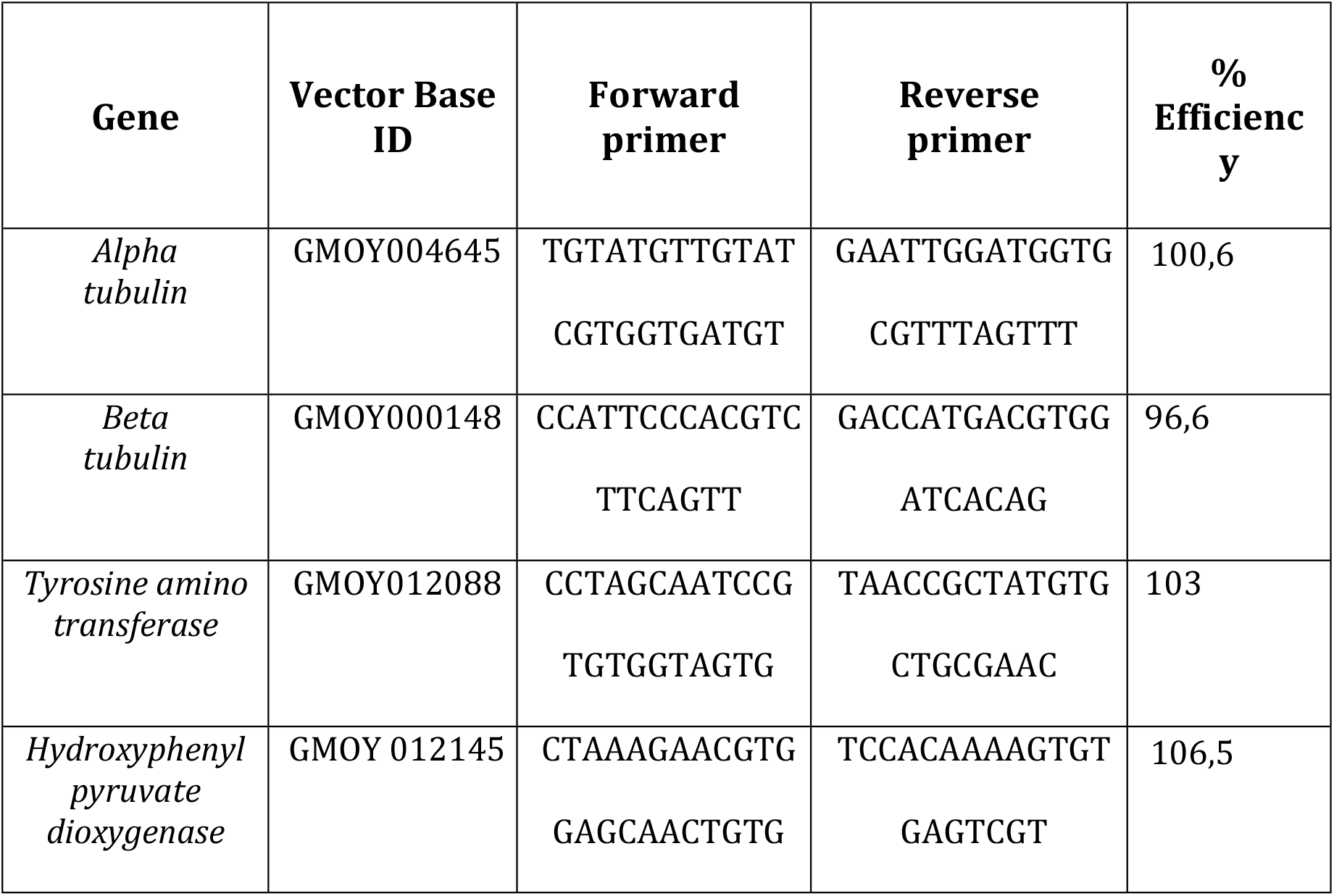
Sequence of the primers used to amplify target genes by Real-Time PCR.

### Oral dosing of mesotrione and nitisinone

Mesotrione (2-[4-(Methylsulfonyl)-2-nitrobenzoyl]cyclohexane-1,3-dione; PESTANAL analytical standard; Sigma-Aldrich) and nitisinone (NTBC, 2-[2-nitro-4-(trifluoromethyl)benzoyl]cyclohexane-1,3-dione; PESTANAL, analytical standard; Sigma-Aldrich) were solubilized in sterile PBS (NaCl 0.15 M, Na-phosphate 10 mM, pH 7.0) to different final concentrations (pH was re-adjusted to 7.0 with 1 M NaOH). One volume of the drugs, or PBS as control, was then mixed with nine volumes of sterile defibrinated horse blood and these spiked bloodmeals were fed to male and female flies. Only fully engorged flies were used. Final concentration in blood for mesotrione were: 2, 1.5, 1, 0.75, 0.5, 0.25, 0.1, 0.05 and 0.01 mg/ml. Final concentration in blood for NTBC were: 0.05, 0.025, 0.01, 0.005, 0.0025, 0.00075, 0.0005, 0.00025, and 0.0001 mg/ml).

### Confocal microscopy

Teneral (<24h old) male tsetse were fed on horse blood (TCS Biosciences) and three days later were offered a second meal containing horse serum with or without 500 ng/mL NTBC. Fifteen hours post-feeding, tsetse were anaesthetized on ice and midguts (with attached proventriculi) were dissected out in ice-cold PBS and immediately fixed in fresh 4% paraformaldehyde for 1 hour at room temperature. Tsetse survival of each group was determined 72 hours post-NTBC administration. Fixed tissues were washed in PBS (and stained with SiR-actin (1:1000 dilution, Cytoskeleton Inc.) for 3 hours at room temperature (RT) and posteriorly incubated in 500 ng/mL DAPI for 10 minutes at RT. Tissues were finally suspended in 1% (w/v) low-melting agarose at ~40°C mixed with Slowfade Diamond oil (Molecular Probes) on a slide. Slides were imaged using a Zeiss LSM-880 confocal laser scanning microscope and tissues were 3D-reconstructed from a series of z-stacks at intervals of 1.3 μm (Zeiss Company).

### Drug treatment of trypanosome-infected flies

Newly emerged (24–48 h. post-eclosion), teneral male *G. m. morsitans* were fed an infected bloodmeal containing *T. brucei* (strain Antat 1.1 90:13)^46^ at 10^6^ parasites per ml of blood. Newly emerged flies are highly susceptible to trypanosome infections. Nine days after infection, when procyclic trypanosomes should be established in the midgut, the flies were fed with blood spiked with NTBC as indicated above.

### Oral dosing of NTBC supplemented with phenylthiourea

*G. m. morsitans* were fed blood spiked with NTBC (0.01 mg/ml) and phenylthiourea (PESTANAL analytical standard; Sigma-Aldrich. 0.1 mg/ml) as indicated above.

### Oral dosing of NTBC in serum

Horse red blood cells were removed by centrifugation at 1000 x *g* for 5 min. The serum was collected, supplemented with mesotrione as indicated above and subsequently fed to *G. m. morsitans*.

### Feeding fructose supplemented with BSA and NTBC

Teneral, male *G. m. morsitans* were sorted into 10 cages and starved for 48 hours. Bovine serum albumin (BSA) fraction V (stock concentration at 200 mg/ml (w/v) in PBS) was serially diluted into sterile 0.1% (w/v) fructose in PBS to create final concentrations of 34, 25, 20, 18, 15, 12, 10, and 5 mg/ml BSA. Control flies were fed with either diluent alone (0.1% fructose) or the diluent spiked with NTBC. All experimental groups were exposed to a NTBC dose of 0.001 mg/ml (lethal when delivered with blood). Flies were offered a meal at 72 hours post-emergence and mortality was tracked daily for one week.

### Administering HPLA to tsetse by feeding or injection

Newly emerged, male *G. m. morsitans* were sorted into 10 cages containing 25 flies/cage. At 48 hours post-emergence, each cage was fed on defibrinated horse blood spiked with distilled water (control) or a serial dilution of DL-p-hydroxyphenyllactic acid (HPLA: Aldrich-Sigma cat no. H3253). All dilutions were made with the same ratio: 25 μl of additive to 1.975 ml of blood. The concentrations of HPLA tested were 0.05, 0.025, 0.01, 0.005, 0.0025, 0.0075, 0.0005, 0.00025, and 0.0001 mg/ml. All dilutions were kept on ice until feeding to avoid chemical degradation. Fresh HPLA stock concentration was 2 mg/ml in sterile water. After 3 days, flies were offered a normal bloodmeal to test if HPLA impaired a second feed. To administer HPLA by injection, one-week old male flies that had taken three previous bloodmeals (to ensure good health and adequate rehydration) were separated into 10 cages. Each group received 2 μl of the selected HPLA concentration via thoracic injection. Stock HPLA was 2 mg/ml and the concentrations injected were 1.0, 0.5, 0.1, 0.05, 0.01, 0.005, 0.001, 0.0005, and 0.0001 mg/ml with sterile water injected as a control. Flies were allowed to recover for 24 hours and then offered a normal bloodmeal. Fly mortality was monitored throughout in both experiments for a week after HPLA administration and no group exceeded 10% mortality.

### *In vivo* oral nitisinone administration to rats

Rats received an oral dose of NTBC by gavage solubilized in sterile PBS. The doses administered were 0.1, 0.2, 0.5, 1.0, and 2.0 mg/kg, with controls receiving an equal volume of PBS. After one and half hours, the rats were anaesthetized with an intraperitoneal injection of 80 μl of 3% ketamine and Xilazin 0.33% diluted in sterile PBS. Neither ketamine nor Xilazin influence tsetse mortality. Groups of male *G. pallidipes* were fed on either PBS or NTBC-treated rats.

### Topical application of mesotrione and NTBC

Mesotrione and NTBC were solubilized in 100% acetone at different concentrations (Mesotrione: 10, 5, 2.5,1, 0.5, and 0.1 mg/ml. NTBC: 1, 0.5, 0.25, 0.125, 0.0625, 0.03175, 0.0156, 0.0078, and 0.0039 mg/ml). One microliter was topically applied to the ventral surface abdomen of cold-anaesthetized *G. m. morsitans*, either immediately before ingesting a blood meal, at different times before or after a bloodmeal. The control flies received 1 μl of 100% acetone. Only fully engorged flies were used.

### Metabolomic analysis

Male *G. m. morsitans* were dissected at different times (0, 5 or 10 hours) after ingesting horse blood spiked with NTBC 0.01 mg/ml or PBS. The flies were chilled for 10 min at 4°C and kept on ice. The legs of 20 flies were severed and collected into 1.5 ml plastic tubes containing 300 μl of 0.9% NaCl and 13.34 μM phenylthiourea (internal standard) in ultrapure water. The samples were vortexed for 1 min, centrifuged for 10 min at 14000 rpm and the supernatant was collected. The supernatants were then filtered (Millipore 0.22 μm syringe filter) to remove cells, such as haemocytes, present in the haemolymph. Filtered samples (150 μl total volume) were transferred to a 1.5 ml tube containing 50 μl of chloroform, 75 μl of methanol and 75 μl of water containing B-methylamino-L-Alanine 53.4 μM as an internal standard for normalization. After centrifugation, the polar and apolar phases were collected and dried in a speed vac concentrator (ISS100. Thermo Scientific). These samples were stored at −80 until analysis. Five samples in total were collected from two independent experiments for each group and time point. See Supplementary Information for details about LC-MS and LC-MS/MS analyses.

### Metabolomic pathway analysis and heat map

The mean relative abundances of all identifiable metabolites for each time-point for both treated and untreated flies were divided by the mean abundance of the metabolite at the corresponding time-point from untreated flies. Results were expressed in fold change values for each individual metabolite, with the untreated group serving as the baseline. Thus, metabolites with abundance fold-changes greater than one were said to be up regulated in flies treated with NTBC as compared to control, whilst metabolites with abundance fold change less than one were said to be down regulated.

The abundance fold-change values were then plotted onto a heatmap using gplot’s heatmap.2 program (version 3.0.1.1, http://cran.r-project.org). The heatmap colour scale was limited from 0 to 4, with additional non-cosmetic options as follows: distfun = function(x) dist(x, “manhattan”), hclustfun = function(x) hclust(x, “centroid”), Colv = FALSE.

Subsequent to this, metabolic pathways were assembled using Pathway Collages from BioCyc^47,48^, focused on pathways relevant to tyrosine degradation, carbon metabolism, and amino acid biosynthesis focused on. Fold-changes from all identified metabolites were overlaid onto this ‘combined’ metabolic pathway, and statistically significant metabolites (as determined by the two-way ANOVA test as described previously) were highlighted. Metabolites that were identified from LC-MS/MS but were not identifiable within the ‘combined’ metabolic pathway, were not mapped. Pathways that did not have any LC-MS/MS-identified metabolites mapped to it were removed from the ‘combined’ pathway.

### *In vitro* comparative metabolism of nitisinone against microsomal preparation from blood-feeding insects

#### Microsome preparation

Mosquito (*Anopheles gambiae* (Kisumu) and *Aedes aegypti* (New Orleans) microsomes were prepared from young female flies (48 to 72 hours old) as described by Kasai *et al*.^49^ and Inceoglu *et al*.^50^ with some modifications. Briefly, preparation of the adult mosquito microsomes (~200 individuals per colony) were done by removing the heads to avoid enzyme inhibition by xanthommatin eye pigments^51^. Mosquitoes were snap-frozen in liquid nitrogen with a sieve ~2 mm mesh size. Small steel spheres were gently mixed with the mosquitoes to fractionate their bodies and to separate the heads, legs, and wings parts from the joint abdominal-thoracic components. The thorax and the abdomen were then washed with 3 ml of pre-chilled 0.1 M KBP pH 7.4 homogenization buffer (HB) containing 1x protease inhibitor (Roche Complete ULTR). The complex was homogenized in 20 ml of HB using 40 ml glass Dounce homogenizer with a loose B pestle (Wheaton Science, Millville, NJ) for 20 strokes. The separation of the homogenate into cytosolic and microsomal extracts was achieved by centrifugation steps at 4°C. The initial centrifugation was performed at 10,000 x *g* for 15 min to remove insoluble debris. A second centrifugation at 200,000 x *g* for 1 h was performed to pellet the microsomes. The microsomal pellet was re-suspended in 2ml of ice-cold suspension buffer (HB containing 20% glycerol). Protein concentration was measured in triplicate by the Bradford method ^52^ with Bio-Rad reagents using bovine serum albumin as a protein standard.

### Tsetse microsomes

Virgin female *G. m. morsitans* (n=174) were maintained on three bloodmeals over the week, and then starved for six days to reduce bloodmeal contaminants during tissue homogenization. The intact thorax and abdomen were removed by a pair of forceps, snap frozen in liquid nitrogen and homogenized as described with mosquito microsomes. Because of the low P450 content in insecticide-susceptible flies and mosquitoes, it was not possible to determine P450 content by the traditional CO-reduced spectral assay^53^, thus protein concentrations were adjusted to 4 mg/ml using 0.1 M Potassium Phosphate buffer (pH 7.4) to normalize for protein content. Recombinant *Anopheles gambaie* CYP6P3 enzyme was used as a positive control for P450s microsomal activities. All microsome preparations, in comparison to CYP6P3, were tested for O-dealkylation activities against the P450’s generic substrate “Diethoxyfluorescein” before the set-up of the NTCB metabolism assay that identifies P450s enzyme activities in microsomal extracts. Briefly, for activity assays, 200μl of reaction media were set up in triplicate containing 0.5 mM NADPH, 5 μM diethoxyfluorescein, 4 mg/ml microsomes or with mixture of 0.1 μM CYP6P3, 0.8 μM cytochrome *b5* and incubated at 30°C for 15 min. The enzyme activity was recorded versus negative control (no NADPH) as 147.93±32, 408.64±45.96, 51.5±2.63 and 372±9.8 relative fluorescence units (RFU) for tsetse, *Ae. aegypti, An. gambiae*, and CYP6P3 respectively. This showed the suitability of microsomes, CYP6P3 and NADPH for running the NTBC comparative metabolism. See Supplementary Information for details about insecticide metabolism experiments.

### NTBC stability under environmental stressors

NTBC was diluted in sucrose 10% to a final concentration of 0.01 mg/ml and stored at either 4°C or 25°C in dark or transparent 1.5 ml polypropylene tubes. One volume of NTBC was then mixed with nine volumes of sterile defibrinated horse blood (final NTBC concentration: 0.001 mg/ml) and these spiked bloodmeals were used to weekly screen against tsetse throughout a five-week period.

Additionally, NTBC was diluted in sucrose 10% (w/v) to a final concentration of 0.01 mg/ml and subjected to ten freeze-thaw cycles. Following temperature stress, one NTBC volume was mixed with nine volumes of sterile defibrinated horse blood (final concentration: 0.001 mg/ml) and these spiked bloodmeals were fed to tsetse. Mortality was monitored over 24 hours.

### Mathematical modelling

A discrete time (1-day time step) compartmental model was constructed to describe the key processes underlying the transmission of *Trypanosoma brucei gambiense*. This parasite has a broad reservoir of host animals, so these were categorised according to whether they were livestock or wildlife. The transmission between three host types in total (humans, livestock and wildlife) and the tsetse fly vector was simulated (see Supplementary Information). The impact of vector biting behaviour was examined by apportioning the split of bites on humans, livestock and wildlife randomly across the full possible range (500 iterations) and, through simulation, determining the frequency of NTBC application required to drive the basic reproduction number (R0) below unity.

### Statistical analysis

Tsetse survival was scored daily post-treatment. Statistical analysis and graphics were performed using Prism 6.0 software (GraphPad Software, San Diego, CA). The data from multiple experiments were combined into a single graph. The log-rank (Kaplan-Meier) test was used to evaluate significant differences in survival between the experimental and control groups. Dose-response curves were done using a plot of non-linear fit for log of inhibitor vs. normalized response (variable slope). LD_50_ and LC_50_ were calculate using Probit analysis (POLO Plus version 2.0). The exact *n* (number of insects) in each experimental replicate used to calculate the statistic in different experiments are detailed in table S1.

## Acknowledgements

This work was partially supported by MRC Confidence in Concept (CiC) awards 2016-17 MC_PC_16052 and 2017-18 MC_PC_17167 (to AA-S and MIP), and BBSRC Anti-VeC award AV/PP0021/1 (to AA-S and LHR). MS was supported by grants from FONCyT (PICT 2017-1015), ANTIVEC (AV/TTKE/0011) and CAPES/FAPERJ (No E-26/ 102.837/2011). PLO was supported by FAPERJ, CAPES and CNPq. SW was supported by LSTM Research Computing Unit. JIM and MSS were supported by The Francis Crick Institute which receives its core funding from Cancer Research UK (FC001999), the UK Medical Research Council (FC001999), and the Wellcome Trust (FC001999). Confocal images were supported by a Wellcome Trust Multi-User Equipment grant (104936/Z/14/Z). We thank Daniel Southern, Keith Steen and Robert Leyland for excellent technical assistance.

## Author contributions

**MS**: conceived the project, designed and performed experiments and wrote the paper. **LRH**: designed and performed experiments and wrote the paper. **AC-S**: performed the experiments with *T. brucei*-infected flies and microscopy analysis. **VOD and DM**: performed *in vivo* experiments. **RJVA and SMB**: performed the experiments with bees. **CR**: performed the phylogenetic analysis and related experiments. **MS, MSS, JM, SQ and SW**: performed metabolomics and analysed the data. **NGE:** performed the experiments related to NTBC stability. **HI and MIP**: performed P450 assays. **LY**: performed the mathematical modelling. **PLO**: conceived the project and provided financial support. **AAS**: conceived the project, designed the experiments, wrote the paper and provided financial support. All authors were involved in proofreading and manuscript correction.

## References

1. Simarro, P., Franco, J., Diarra, A. & Jannin, J. Epidemiology of human African trypanosomiasis. Clin. Epidemiol. 257 (2014) doi:10.2147/CLEP.S39728.

2. WHO | Trypanosomiasis, human African (sleeping sickness). WHO (2016).

3. WHO. Strategic review of traps and targets for tsetse and african trypanosomiasis control. World Heal. Rep. 1–58 (2004).

4. Rayaisse, J.-B. et al. Standardizing visual control devices for tsetse flies: west africanspecies *Glossina tachinoides, G. palpalis gambiensis* and *G. morsitans submorsitans*. PLoS Negl. Trop. Dis. 6, e1491 (2012).

5. Rayaisse, J. B. et al. Prospects for the development of odour baits to control the tsetse flies *Glossina tachinoides* and *G. palpalis*. PLoS Negl. Trop. Dis. 4, e632 (2010).

6. Ndeledje, N. et al. Treating cattle to protect people? Impact of footbath insecticide treatment on tsetse density in Chad. PLoS One 8, e67580 (2013).

7. Torr, S. J., Maudlin, I. & Vale, G. A. Less is more: restricted application of insecticide to cattle to improve the cost and efficacy of tsetse control. Med. Vet. Entomol. 21, 53–64 (2007).

8. Simo, G. & Rayaisse, J. B. Challenges facing the elimination of sleeping sickness in west and central Africa: sustainable control of animal trypanosomiasis as an indispensable approach to achieve the goal. Parasit. Vectors 8, 640 (2015).

9. M. J. Lehane. The Biology of Blood-Sucking in Insects. Psychological Science vol. 2nd Edn (Liverpool School of Tropical Medicine, 2005).

10. Sterkel, M., Oliveira, J. H. M., Bottino-Rojas, V., Paiva-Silva, G. O. & Oliveira, P. L. The dose makes the poison: nutritional overload determines the life traits of blood-feeding arthropods. TrendsParasitol. 33, 633–644 (2017).

11. Sterkel, M. et al. Tyrosine detoxification is an essential trait in the life history of blood-feeding arthropods. Curr. Biol. 26, 2188–2193 (2016).

12. Holme, E. & Lindstedt, S. Tyrosinaemia type I and NTBC (2-(2-nitro-4-trifluoromethylbenzoyl)-1,3-cyclohexanedione). J. Inherit. Metab. Dis. 21, 507–17 (1998).

13. Lock, E., Ranganath, L. R. & Timmis, O. The role of nitisinone in tyrosine pathway disorders. Curr. Rheumatol. Rep. 16, 457 (2014).

14. Genome sequence of the pea aphid *Acyrthosiphon pisum*. PLoS Biol. 8, e1000313 (2010).

15. Nowicki, C. & Cazzulo, J. J. Aromatic amino acid catabolism in trypanosomatids. Comp. Biochem. Physiol. Part A Mol. Integr. Physiol. 151, 381–390 (2008).

16. Watanabe, J. et al. Genome sequence of the tsetse fly *(Glossina morsitans):* vector of african trypanosomiasis. Science (80-.). 344, 380–386 (2014).

17. Hall, M. G., Wilks, M. F., Provan, W. M., Eksborg, S. & Lumholtz, B. Pharmacokinetics and pharmacodynamics of NTBC (2-(2-nitro-4-fluoromethylbenzoyl)-1,3-cyclohexanedione) and mesotrione, inhibitors of 4-hydroxyphenyl pyruvate dioxygenase (HPPD) following a single dose to healthy male volunteers. Br. J. Clin. Pharmacol. 52, 169–177 (2001).

18. Lindstedt, S. Treatment of hereditary tyrosinaemia type I by inhibition of 4-hydroxyphenylpyruvate dioxygenase. Lancet 340, 813–817 (1992).

19. Lock, E., Ranganath, L. R. & Timmis, O. The role of nitisinone in tyrosine pathway disorders. Curr. Rheumatol. Rep. 16, 1–8 (2014).

20. González-Santoyo, I. & Córdoba-Aguilar, A. Phenoloxidase: a key component of the insect immune system. Entomol. Exp. Appl. 142, 1–16 (2012).

21. Ryazanova, A. D., Alekseev, A. A. & Slepneva, I. A. The phenylthiourea is a competitive inhibitor of the enzymatic oxidation of DOPA by phenoloxidase. J. Enzyme Inhib. Med. Chem. 27, 78–83 (2012).

22. Graça-Souza, A. V et al. Adaptations against heme toxicity in blood-feeding arthropods. Insect Biochem. Mol. Biol. 36, 322–35 (2006).

23. Riond, B., Wenger-Riggenbach, B., Hofmann-Lehmann, R. & Lutz, H. Serum protein concentrations from clinically healthy horses determined by agarose gel electrophoresis. Vet. Clin. Pathol. 38, 73–77 (2009).

24. Huledal, G. et al. Non randomized study on the potential of nitisinone to inhibit cytochrome P450 2C9, 2D6, 2E1 and the organic anion transporters OAT1 and OAT3 in healthy volunteers. Eur. Jou. Clin. Phar. 75, 313–320 (2019)

25. Jason N. Neat, Andrea Wolff, Faraz Kazmi, Pilar Prentiss, David Buckley, Elizabeth M. Wilson, John Bial, Cedo M. Bagi, M. G. and A. P. *In vitro* inhibition and induction of human liver cytochrome P450 enzymes by NTBC and its metabolism in human liver microsomes. Drug Metab. Rev. 42, (2010).

26. Müller, P. et al. Field-caught permethrin-resistant *Anopheles gambiae* overexpress CYP6P3, a P450 that metabolises pyrethroids. PLoS Genet. 4, e1000286 (2008).

27. Barchanska, H., Rola, R., Szczepankiewicz, W. & Mrachacz, M. LC-MS/MS study of the degradation processes of nitisinone and its by-products. J. Pharm. Biomed. Anal. 171, 15–21 (2019).

28. Klein, A.-M., Boreux, V., Fornoff, F., Mupepele, A.-C. & Pufal, G. Relevance of wild and managed bees for human well-being. Curr. Opin. Insect Sci. 26, 82–88 (2018).

29. Mužinić, V. & Želježić, D. Non-target toxicity of novel insecticides. Arh. Hig. Rada Toksikol. 69, 86–102 (2018).

30. Pooda, S. H., Mouline, K., De Meeûs, T., Bengaly, Z. & Solano, P. Decrease in survival and fecundity of *Glossina palpalis gambiensis* vanderplank 1949 (Diptera: Glossinidae) fed on cattle treated with single doses of ivermectin. Parasit. Vectors 6, 165 (2013).

31. Distelmans, W., D’Haeseleer, F. & Mortelmans, J. Efficacy of systemic administration of ivermectin against tsetse flies. Ann. Soc. Belg. Med. Trop. (1920). 63, 119–25 (1983).

32. Van den Abbeele, J., Van den Bossche, P., Mortelmans, J. & Decleir, W. Effect of ivermectin and isometamidium chloride on *Glossina palpalis palpalis* (Diptera: Glossinidae). Ann. Soc. Belg. Med. Trop. (1920). 68, 53–9 (1988).

33. Van Den Bossche, P. & Geerts, S. The effects on longevity and fecundity of *Glossina tachinoides* after feeding on pigs treated with ivermectin. Ann. Soc. Belg. Med. Trop. (1920). 68, 133–139 (1988).

34. European Medicines Agency-Orfadin, INN-Nitisinone. 1, 1–102 (2004).

35. DrugBank. https://www.drugbank.ca/.

36. Sterkel, M., Ons, S. & Oliveira, P. L. DOPA decarboxylase is essential for cuticle tanning in *Rhodnius prolixus* (Hemiptera: Reduviidae), affecting ecdysis, survival and reproduction. Insect Biochem. Mol. Biol. 108, 24–31 (2019).

37. Macagnan, N. et al. Toxicity of cypermethrin and deltamethrin insecticides on embryos and larvae of *Physalaemus gracilis* (Anura: Leptodactylidae). Environ. Sci. Pollut. Res. 24, 20699–20704 (2017).

38. Gurevitz, J. M. et al. Unexpected failures to control Chagas Disease vectors with pyrethroid spraying in northern Argentina. J. Med. Entomol. 49, 1379–86 (2012).

39. Toé, K. H. et al. Increased pyrethroid resistance in malaria vectors and decreased bed net effectiveness, Burkina Faso. Emerg. Infect. Dis. 20, 1691–1696 (2014).

40. Strode, C., Donegan, S., Garner, P., Enayati, A. A. & Hemingway, J. The impact of pyrethroid resistance on the efficacy of insecticide-treated bed nets against african anopheline mosquitoes: systematic review and meta-Analysis. PLoS Med. 11, e1001619 (2014).

41. Dereeper, A. et al. Phylogeny.fr: robust phylogenetic analysis for the non-specialist. Nucleic Acids Res. 36, W465–W469 (2008).

42. Dereeper, A., Audic, S., Claverie, J.-M. & Blanc, G. BLAST-EXPLORER helps you building datasets for phylogenetic analysis. BMC Evol. Biol. 10, 8 (2010).

43. Guindon, S. et al. New Algorithms and methods to estimate maximum-likelihood phylogenies: assessing the performance of PhyML 3.0. Syst. Biol. 59, 307–321 (2010).

44. Walshe, D. P., Lehane, S. M., Lehane, M. J. & Haines, L. R. Prolonged gene knockdown in the tsetse fly Glossina by feeding double stranded RNA. Insect Mol. Biol. 18, 11–19 (2009).

45. Livak, K. J. & Schmittgen, T. D. Analysis of relative gene expression data using real-time quantitative PCR and the 2^-ΔΔCT^ Method. Methods 25, 402–408 (2001).

46. MacGregor, P., Rojas, F., Dean, S. & Matthews, K. R. Stable transformation of pleomorphic bloodstream form *Trypanosoma brucei*. Mol. Biochem. Parasitol. 190, 60–62 (2013).

47. Paley, S., O’Maille, P. E., Weaver, D. & Karp, P. D. Pathway collages: personalized multi-pathway diagrams. BMC Bioinformatics 17, 529 (2016).

48. Karp, P. D. et al. Pathway Tools version 19.0 update: software for pathway/genome informatics and systems biology. Brief. Bioinform. 17, 877–890 (2016).

49. Kasai, S. et al. Mechanisms of pyrethroid resistance in the dengue mosquito vector, *Aedes aegypti:* target site insensitivity, penetration, and metabolism. PLoS Negl. Trop. Dis. 8, e2948 (2014).

50. Inceoglu, A. B. et al. A rapid luminescent assay for measuring cytochrome P450 activity in individual larval *Culex pipiens* complex mosquitoes (Diptera: Culicidae). J. Med. Entomol. 46, 83–92 (2009).

51. Schonbrod, R. D.; Terriere, L. C. Inhibition of housefly microsomal epoxidase by the eye pigment, xanthommatin. Pestic. Biochem. Physiol. 1, 409–417 (1971).

52. Bradford, M. A rapid and sensitive method for the quantitation of microgram quantities of protein utilizing the principle of protein-dye binding. Anal. Biochem. 72, 248–254 (1976).

53. Omura, T.; Sato, R. The carbon monoxide-binding pigment of liver microsomes. Solubilization, purification, and properties. J. Biol. Chem 239, 2379–85 (1964).

